# Heterogeneous Nuclear Ribonucleoprotein A1 as a Key Regulator in Pulmonary Arterial Hypertension Development

**DOI:** 10.64898/2026.01.05.697826

**Authors:** Yoshihito Morimoto, Yoshiaki Sato, Katsuhiro Kato, Hidenori Yamamoto, Kentaro Suzuki, Kiyotaka Go, Yoshie Fukasawa, Naoki Ohashi, Yoshiyuki Takahashi, Taichi Kato

**Affiliations:** Department of Pediatrics, Nagoya University Graduate School of Medicine, Nagoya, Aichi, Japan; Department of Pediatric Cardiology, National Cerebral and Cardiovascular Center, Suita, Osaka, Japan; Division of Neonatology, Center for Maternal-Neonatal Care, Nagoya University Hospital, Nagoya, Aichi, Japan; Department of Cardiology, Nagoya University Graduate School of Medicine, Nagoya, Aichi, Japan

**Keywords:** plexiform lesion formation, pulmonary arterial hypertension, unobstructed pulmonary arteries with medial hypertrophy, differentially expressed proteins, rat model

## Abstract

**Background:** Although plexiform lesion (PL) formation in severe pulmonary arterial hypertension (PAH) is a therapeutic target, the mechanisms underlying their formation have not been fully elucidated. This study aimed to identify candidate proteins involved in PL formation by examining differentially expressed proteins (DEPs) in PAH lesions.

**Methods:** Proteomic remodeling was assessed before and after the formation of PLs in a SU5416 combined with hypoxia (SuHx) rat model of severe PAH using laser-capture microdissection coupled with mass spectrometry. Unobstructed pulmonary arteries with medial hypertrophy (UMHPAs) and PLs from SuHx rats were subjected to qualitative and quantitative proteomics, revealing DEPs between these structures.

**Results:** We identified 718 proteins with 58 DEPs, of which 31 were upregulated in UMHPAs and 27 were upregulated in PLs. Immunostaining confirmed that DEPs detected in our proteomic analysis were differentially expressed between UMHPAs and PLs. Among them, we focused on heterogeneous nuclear ribonucleoprotein A1 (hnRNPA1) as a candidate protein that may be strongly associated with PL formation because of its strong association with cell proliferation. siRNA knockdown of hnRNPA1 in hypoxia-treated pulmonary artery smooth muscle cells reduced pyruvate kinase M2 expression and significantly decreased proliferative capacity. Treatment of SuHx rats with hnRNPA1 inhibitors also suppressed pulmonary artery remodeling in PAH and PL formation.

**Conclusions:** Several DEPs associated with PL formation but with unclear relevance to pulmonary artery remodeling in PAH were discovered. Among these, hnRNPA1, which was not detected in transcriptome analysis, may be important in PL formation.

**What are the Clinical Implications?:** The prognosis for pulmonary arterial hypertension (PAH) with plexiform lesions (PLs) is poor. Although PLs are therapeutic targets, the mechanisms of PL formation have not been fully elucidated, and there have been no direct examinations of proteomic remodeling before and after their formation. We identified several differentially expressed proteins (DEPs) that may be strongly associated with PL formation. Of these, heterogeneous nuclear ribonucleoprotein A1 (hnRNPA1) was examined as a candidate protein involved in PA remodeling in PAH and PLs. Inhibition of hnRNPA1 in a severe PAH model not only suppressed pulmonary artery remodeling in PAH but also reduced PL formation. Several DEPs involved in pulmonary artery remodeling in PAH may be of significant value in the search for new therapeutic targets.

## INTRODUCTION

Pulmonary arterial hypertension (PAH), a life-threatening disease, starts with thickening of the tunica media and progresses to intimal proliferation and generation of obstructive lesions. Severe PAH eventually leads to plexiform lesions (PLs) with necrosis of the tunica media and disruption of the vascular structure.^1^ Conventional drugs alleviate lumen narrowing caused by tunica media thickening through pulmonary vasodilation. New promising drugs such as sotatercept (an activin signaling inhibitor) have become available, but the prognosis for severe PAH that forms PLs remains poor.^2, 3^ Therefore, it is imperative to identify therapeutic targets to inhibit PL formation. However, the mechanisms underlying PL formation remain unclear. One reason is that PLs coexist with other tissues in the lungs, presenting challenges in analyzing PLs in isolation.

Laser-capture microdissection coupled with mass spectrometry (LCM–MS) is an efficient method for the systematic identification and quantification of proteins in tissue samples. This method can accurately detect regional tissue differences owing to its ability to maintain the integrity of data related to protein localization and spatial relationships.^4–6^ Although proteomic analyses have been performed on whole lungs from PAH patients, containing several other tissues, few LCM–MS procedures have been performed on pulmonary arteries (PAs), and there have been no reports on PLs.^7–9^ This may be because PAs are luminal structures, and it is not easy to obtain the necessary amount of tissue for LCM–MS. Similar research methods involving microRNA transcriptomes after LCM for PLs have been reported^2, 10^; however, there is a substantial discrepancy of approximately 50% between transcriptome and proteome analysis results, potentially resulting in divergent conclusions.

Although PLs are therapeutic targets, there have been no direct examinations of proteomic remodeling before and after their formation. Here, we examined proteomic remodeling using LCM–MS before and after PL formation in a rat model of severe PAH involving injection of SU5416, a selective inhibitor of vascular endothelial growth factor receptors, combined with hypoxia (SuHx).^11^ We performed LCM–MS using formalin-fixed paraffin-embedded (FFPE) sections. Based on proteome analysis results, we performed a functional analysis of heterogeneous nuclear ribonucleoprotein A1 (hnRNPA1) as a candidate protein involved in PA remodeling in PAH and PLs.

## MATERIALS AND METHODS

Please see the Supplemental Methods and the Major Resources Table in the Supplemental Material for details of some experimental procedures.

### Data Availability

The data that support the findings of this study are available from the corresponding author upon reasonable request.

### Generation of the SuHx Rat Model

All animals received humane care, and all experimental procedures were approved by the Animal Care and Use Committee of Nagoya University (M240320-001 and M240315-001). In addition, all procedures conformed to the guidelines from Directive 2010/63/EU of the European Parliament on the protection of animals used for scientific purposes. We generated a SuHx rat model as described in the Supplemental Methods.^11^

### Hemodynamic Measurements and Sample Collection

Hemodynamic measurement and sample collection were performed as described in the Supplemental Methods.

### Hematoxylin–Eosin and Toluidine Blue Staining

Staining of samples was performed as described in the Supplemental Methods.

### Histological Measurements

The histological measurement method was adopted from the procedure described by Shinohara et al.^12^ and the slides were analyzed in a blinded manner (Supplemental Methods). In addition, all small arteries (outer diameter <50 μm) per lung section were assessed for occlusive lesions at 40× magnification. A vessel in which the lumen was partially (50%) or fully obstructed was defined as an occlusive lesion (Grade 2).^13^ Quantitative analysis was performed to determine the ratio of occlusive lesions among all the small PAs per lung section. The types of vessel occlusion were defined by the pattern of α-smooth muscle actin (αSMA)-positive cell distribution.^14^

### Laser-Capture Microdissection

Regions of unobstructed pulmonary arteries with medial hypertrophy (UMHPAs) and PLs were isolated using an LMD7000 system (Leica, Germany) at 20× magnification. Concentric laminar endometrial fibrosis (onion skin lesions) associated with PLs, was included in PLs. Each tissue was collected in a 2.5 mm^2^ area (n = 5 per group) as reported earlier^7, 8, 15^ (Supplemental Methods). Only UMHPAs from the 8w model and PLs from the 13w model were collected (Figure S1).

### Tissue Digestion for Mass Spectrometry

This method was adapted from a modified version of the procedure described by Drummond et al.^15, 16^ For detailed procedures, please refer to the Supplemental Methods.

### Mass Spectrometry Data Analysis

The peptides were analyzed via liquid chromatography–tandem mass spectrometry (LC–MS/MS), as detailed in the Supplemental Methods.

### Immunohistochemistry

Immunostaining using the Ventana Discovery Ultra Immunostainer (Roche, Switzerland) was performed to validate the proteomic results and expression site of DEPs. All protocols followed the manufacturer’s recommendations. The localization and intensity of immunoreactivity were determined by an investigator blinded to the experimental and control groups. Primary antibodies specific to the proteins listed in the Major Resources Table were used.

### Pathway Analysis

For each protein upregulated in UMHPAs and PLs, pathway and process enrichment analyses were performed using Metascape (https://metascape.org/gp/index.html). Only terms with *P* < 0.01, a minimum count of 3, and an enrichment factor (ratio between the observed counts and counts expected by chance) >1.5 were collected and grouped into clusters based on membership similarities.

### Serum hnRNPA1 Concentration Measurement

Among the DEPs, we focused on elevated hnRNPA1 and examined the role of PAH remodeling of PA, based on the literature search. Before functional analysis, serum hnRNPA1 concentration was determined using a rat hnRNPA1 ELISA kit (Cat# abx353728, Abbexa).

### Immunofluorescence to Investigate Localization of hnRNPA1

We performed immunofluorescence on PLs to validate the cell types exhibiting elevated levels of hnRNPA1. Samples were stained as described in the Supplemental Methods.

### Cell Culture and Characterization

Control rat pulmonary artery smooth muscle cells (rPASMCs) were purchased from Cell Applications, USA (Cat# CAR35205a) and cultured in Dulbecco’s modified Eagle’s medium containing 10% fetal bovine serum and 1% penicillin–streptomycin in a humidified incubator at 37 ℃ with 5% CO_2_. Passages 5–8 were used for all experiments. The cells were exposed to hypoxic conditions (5% O_2_) to simulate the effects of PAH.^18^ Hypoxia was induced in an incubator (Cat# MCO-5MUV-PJ, PHC Holdings, Japan) by introducing nitrogen gas, and the oxygen concentration was monitored continuously using an oxygen sensor.

### Small Interfering RNA (siRNA) Transfection

Knockdown experiments were conducted on hnRNPA1 in rPASMCs, as detailed in the Supplemental Methods.

### hnRNPA1 Inhibition in rPASMCs

We examined the role of hnRNPA1 using tetracaine hydrochloride (tetracaine), a known hnRNPA1 inhibitor, because it is inexpensive and drug doses in an experimental melanoma model have been reported.^19^ Experiments were first conducted to ascertain whether tetracaine suppressed hnRNPA1 expression in hypoxia-treated rPASMCs. Briefly, hypoxia-treated rPASMCs were pre-seeded at 1 × 10^4^ cells/well in a 96-well plate for evaluation of cell proliferation or at 2 × 10^5^ cells/well in a 24-well plate for western blotting. Prior to experiments, hypoxia-treated rPASMCs were subjected to 24-h serum starvation to induce quiescence and then treated with 0, 50, 100, 200, or 400 μM tetracaine. Cell proliferation was evaluated at 0, 24, 48, and 72 h and western blotting after 24 h.

### Cell Proliferation Assay

Cell proliferation was measured using a cell counting kit-8 (CCK-8) assay (Cat#:CK04-01, Dojindo, Japan) according to the manufacturer’s instructions (Supplemental Methods).

### Immunofluorescence Analysis for Cultured rPASMCs

Immunofluorescence on cultured rPASMCs was performed as described in the Supplemental Methods.

### Protein Extraction from Cultured rPASMCs

Proteins were extracted from rPASMCs using the Minute Total Protein Extraction Kit for Animal Cultured Cells and Tissues (Cat# SD-001, Invent Biotechnologies, USA), according to the manufacturer’s instructions.

### Sample Preparation for Automatic Capillary Western Blotting

Western blotting was performed using an automatic capillary western blotting device (Simple Western; Jess, ProteinSimple, USA) (Supplemental Methods).

### Inhibition of hnRNPA1 in SuHx Rats with Tetracaine

Once we confirmed that tetracaine inhibited hnRNPA1 in hypoxia-treated rPASMCs, we evaluated the effect of tetracaine in SuHx rats. Adult male Sprague–Dawley rats (n = 12) were injected subcutaneously with SU5416 (20 mg/kg) and exposed to hypobaric hypoxia (0.5 atm) for 3 weeks. Thereafter, SuHx rats were returned to normoxia (21% O_2_) for an additional 10 weeks. Eight weeks after SU5416 administration, when PL formation was in progress,^11^ tetracaine (10 mg/kg; based on drug doses used previously)^16^ was injected intraperitoneally into SuHx rats (treated group; n = 6) once a week (Figure S2). Rats in the non-treated, normal control group (n = 6) were injected intraperitoneally with an equal volume of PBS and maintained in ambient air.

### hnRNPA1 Staining Analysis

We used ImageJ to evaluate hnRNPA1 staining and count the number of cell nuclei. In this application, diaminobenzidene and hematoxylin staining were removed and only hnRNPA1 staining was detected. We measured mean gray values of hnRNPA1 staining as optical density, in order to quantify staining levels.

### Statistical Analysis

Data are presented as the mean ± SE for continuous variables. One-way analysis of variance was used to compare variables among three or more groups, followed by Tukey’s multiple comparison test. When two groups were compared, the Wilcoxon signed-rank test was used to assess significant differences. Results with *P* < 0.05 were considered statistically significant. JMP 13 Pro software (SAS Institute, Cary, North Carolina, USA) was used for all analyses.

## RESULTS

### Hemodynamics and Histopathology of SuHx Rats

Relative to the level in control rats (29.6 ± 1.9 mmHg), right ventricular systolic pressure (RVSP) increased in rats in both 8-week (8w; 105.6 ± 12.5 mmHg, *P* < 0.01) and 13-week (13w; 119.4 ± 11.5 mmHg, *P <* 0.01) models (Figure 1A). Relative to the level in control rats (0.23 ± 0.01), Fulton’s index increased in rats in both 8w (0.55 ± 0.01, *P* < 0.01) and 13w (0.60 ± 0.02, *P <* 0.01) models (Figure 1B).

**Figure 1.**
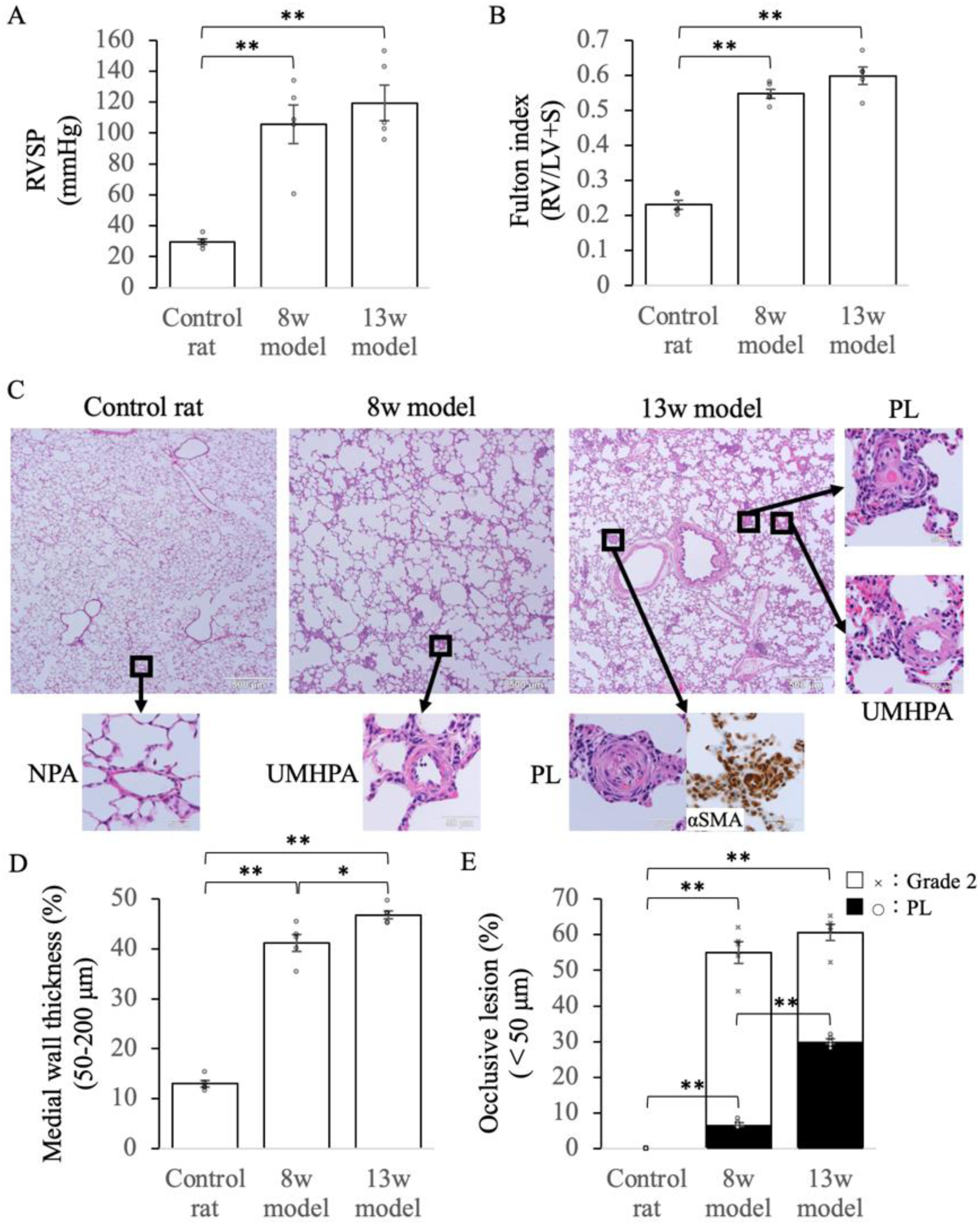
PAH progression in the SuHx rat model. **A**, Right ventricular systolic pressure (RVSP). **B**, Fulton’s index, weight ratio of the right ventricle to the left ventricle + septum (RV/LV+S). **C**, Hematoxylin–eosin staining showing the pathological progression in SuHx rats (scale bar = 500 μm; inset, scale bar = 50 μm). **D**, Percentage of medial wall thickness. **E**, Percentage of occlusive lesions among the vessels. \**P* < 0.05; \*\**P* < 0.01. Statistical analysis was performed using the Wilcoxon signed-rank test. Values are the mean ± SE, n = 5 per group. αSMA, α-smooth muscle actin; PL, plexiform lesion; SuHx, SU5416 combined with hypoxia; 8w, 8-week; 13w, 13-week; NPA, normal pulmonary artery; PL, plexiform lesion; UMHPA, unobstructed pulmonary artery with medial hypertrophy.

Figure 1C–1E shows the pathological progression in SuHx rats. The medial wall thickness of the control rats was thin (13.0 ± 0.7%), and no PLs (0.0 ± 0.0%) were observed. In the 8w model, there were few PLs (6.8 ± 0.6%) but several UMHPAs with thickened medial wall thickness (41.2 ± 1.7%, *P* < 0.01), although not completely occluded; however, in the 13w model, many PLs (30.0 ± 0.7%, *P* < 0.01) were present.

### LCM–MS

We attempted to extract normal PA tissue from control rats but could not obtain sufficient amounts for LCM–MS. Therefore, PLs in the 13w model were compared with UMHPAs during remodeling in the 8w model (Figure S1). LCM–MS identified 718 proteins, with 58 DEPs among these regions. Of these, 31 DEPs were upregulated in UMHPAs during remodeling, and 27 DEPs were upregulated in PLs. Alpha-cardiac muscle 1 was the most upregulated protein in UMHPAs, followed by TAGLN, histone H2A.Z, and ANXA3 (Table S1). In PLs, TMSB4 was the most upregulated protein, followed by MGP, CALML3, and C4 (Table S2).

### Immunohistochemistry

To validate the proteome results and explore the cellular location of DEPs, we performed immunostaining of the top DEPs that were altered according to LCM–MS and visually confirmed the predominant changes (Figure 2). The top DEPs were less expressed in normal PAs compared with those in UMHPAs and PLs. Many of the upregulated DEPs in UMHPAs were proteins expressed on cells lining the lumen of PAs. Upregulated DEPs, including hnRNPA1, were highly expressed in PLs compared with those in normal PAs and UMHPAs.

**Figure 2.**
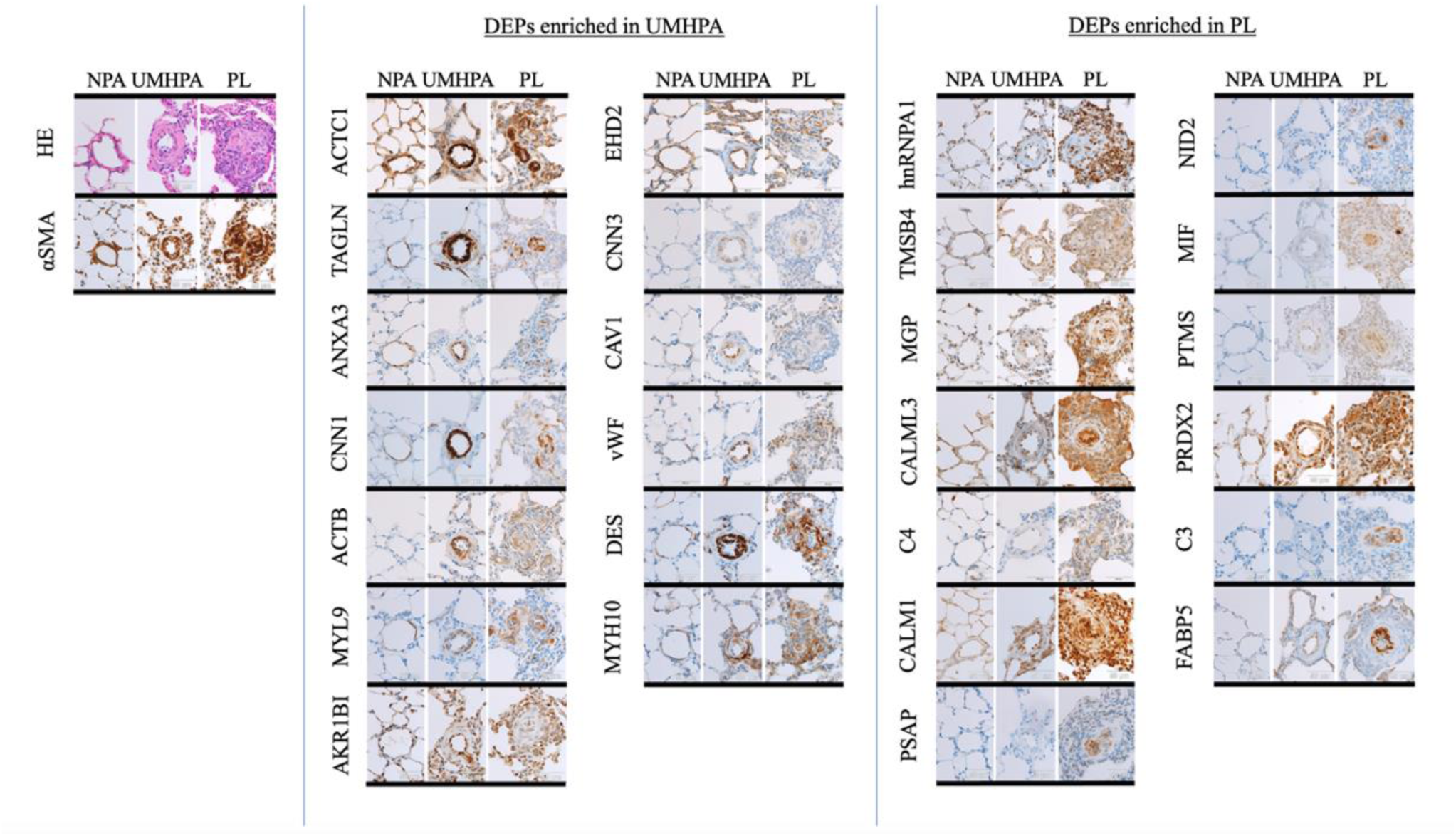
Hematoxylin–eosin staining and immunohistochemistry of DEPs in normal PAs, UMHPAs, and PLs. Hematoxylin–eosin staining results of normal PAs, UMHPAs, and PLs and immunohistochemistry results of αSMA and each DEP are shown (scale bar = 50 μm). ACTB, β-actin; ACTC1, actin alpha-cardiac muscle 1; AKR1B1, aldo-keto reductase family 1 member B; αSMA, α-smooth muscle actin; ANXA3, annexin A3; CALM1, calmodulin 1; CALML3, calmodulin-like 3; CAV1, caveolin 1; CNN1, calponin 1; CNN3, calponin 3; C3, complement C3; C4, complement C4; DEP, differentially expressed protein; DES, desmin; EHD2, EH domain-containing 2; FABP5, fatty acid-binding protein 5; HE, hematoxylin–eosin; hnRNPA1, heterogeneous nuclear ribonucleoprotein A1; MGP, matrix Gla protein; MIF, macrophage migration inhibitory factor; MYH10, myosin heavy chain 10; MYL9, myosin light chain 9; NID2, nidogen 2; NPA, normal pulmonary artery; PL, plexiform lesion; PRDX2, peroxiredoxin 2; PSAP, prosaposin; PTMS, parathymosin; TAGLN, transgelin; TMSB4, thymosin beta 4; UMHPA, unobstructed pulmonary artery with medial hypertrophy; vWF, von Willebrand factor.

### Pathway Analysis

DEPs in UMHPAs and PLs were subjected to pathway and process enrichment analyses (Figure S3). The top three protein clusters upregulated in UMHPAs consisted of actin cytoskeleton organization (GO:0030036), focal adhesion (rno04510), and cell–extracellular matrix interactions (R-RNO-446353), whereas those in PLs consisted of innate immune system (R-RNO-168249), pertussis (rno05133), and G2/M checkpoints (R-RNO-69481).

### Candidate Protein Involved in PA Remodeling in PAH and PLs

From a literature search, we determined that hnRNPA1, which shows elevated levels in PLs, might be a candidate protein involved in PA remodeling in PAH and PLs because it is an RNA-binding protein (RBP) and because of the association of RBPs with PA remodeling in PAH.^20^ Therefore, we performed a functional analysis of hnRNPA1.

The variability of serum hnRNPA1 in SuHx rats was assessed before functional analysis of hnRNPA1. Relative to the level in control rats (23.7 ± 2.3 ng/mL), serum hnRNPA1 concentration increased in the 8w (70.7 ± 12.2 ng/mL, *P* < 0.01) and 13w (63.1 ± 14.3 ng/mL, *P <* 0.01) models (Figure S4). This indicates that hnRNPA1 levels are significantly elevated both in the PAH lesions and sera of SuHx rats.

### Investigation of Localization of hnRNPA1 via Immunofluorescence

Figure S5 shows the localization of hnRNPA1. Compared to UMHPAs, immunofluorescence also clearly showed enhanced hnRNPA1 expression in PLs. In UMHPAs, some αSMA-positive cells (derived from PA-derived smooth muscle cells; PASMCs) were hnRNPA1-positive, but PECAM-1-positive cells (pulmonary vascular endothelial cell-derived cells) were hnRNPA1-negative. In PLs, αSMA-positive cells and LCA-positive cells (immune cell-derived cells) showed strong positivity for hnRNPA1, whereas PECAM-1-positive cells were hnRNPA1-negative. This finding indicates that hnRNPA1 expression is enhanced in the cellular composition of PLs, specifically in PASMCs and immune cell-derived cells.

### siRNA Inhibition of hnRNPA1 in Normal Mature and Hypoxia-Treated Rat PASMCs

To analyze the function of hnRNPA1, which may be involved in PA remodeling in PAH and PLs, rat PASMC knockdown experiments were performed using siRNAs because αSMA-positive cells were hnRNPA1-positive following immunofluorescence. Figure 3A illustrates the proliferative capacity of rPASMCs (measured using the CCK-8 assay). Figure 3B shows rPSAMCs stained with Hoechst dye 72 h after siRNA transfection, whereas Figure 3C shows the number of rPASMCs adhering to the bottom of 96-well plates 72 h after siRNA transfection. Figure 3D and 3E show the proliferative capacity (assessed by Ki-67-positive cell content) and apoptosis (assessed by cleaved caspase-3-positive cell content) of rPASMCs, respectively, demonstrating that transfection of normal rPASMCs with hnRNPA1 siRNA reduced their proliferative capacity and increased apoptosis. We used hypoxia-treated rPASMCs to simulate arterial muscle cells in pulmonary vascular lesions. Hypoxia-treated rPASMCs exhibited higher proliferative capacity and suppressed apoptosis compared with normal rPASMCs, whereas transfection with hnRNPA1 siRNA strongly suppressed proliferative capacity and increased apoptosis.

**Figure 3.**
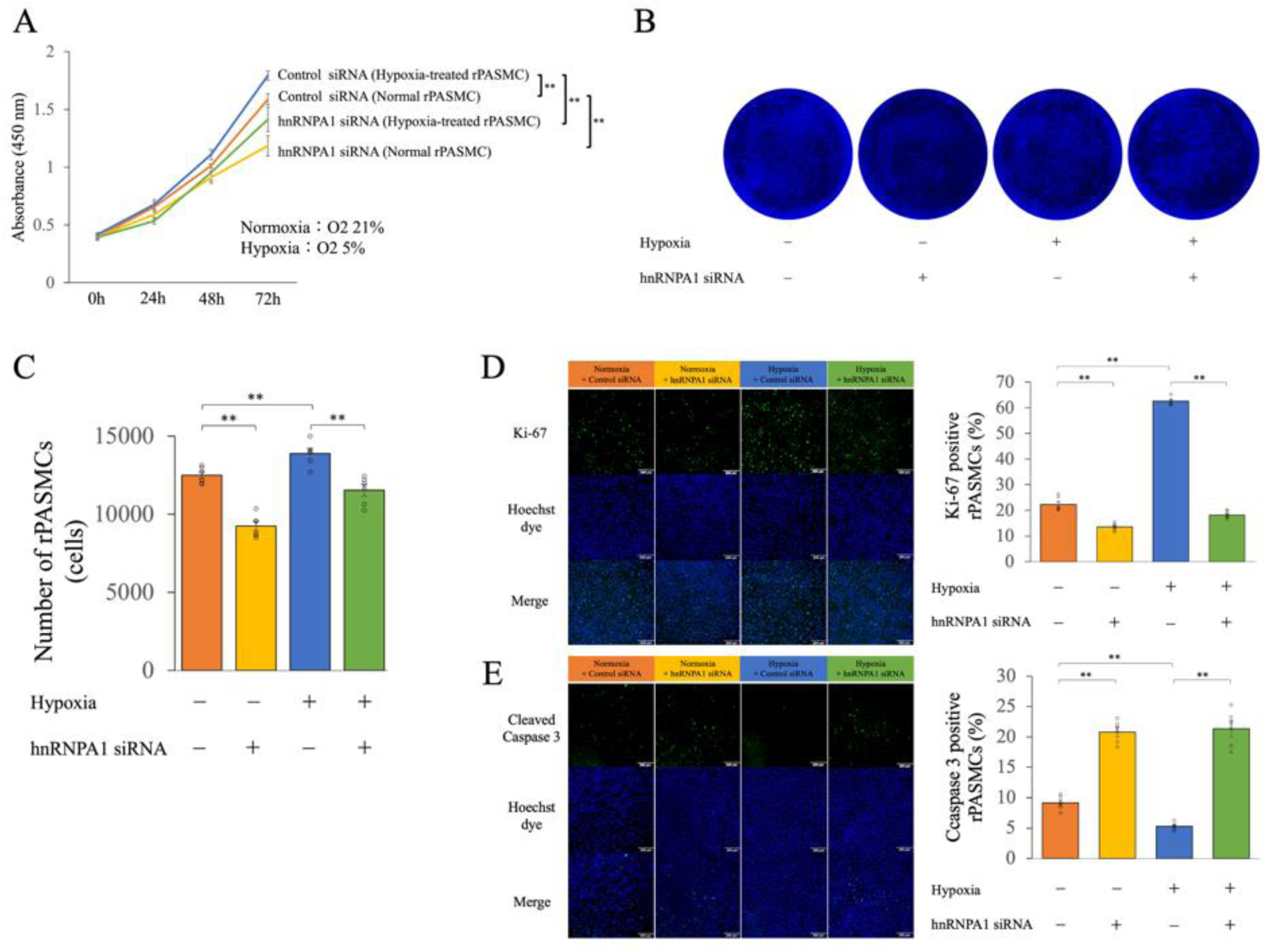
Effect of hnRNPA1 siRNA on proliferative capacity and apoptosis in normal rPASMCs and hypoxia-treated rPASMCs. **A**, Proliferative capacity of rPASMCs in the hnRNPA1 siRNA-treated group compared with that in the negative control siRNA-treated group, as measured using the CCK-8 assay. **B**, rPSAMCs stained with Hoechst dye 72 h after siRNA transfection. **C**, Number of rPASMCs adhering to the bottom of 96-well plates 72 h after siRNA transfection. **D**, Proliferative capacity of rPASMCs in the hnRNPA1 siRNA-treated group compared with the negative control siRNA-treated group via Ki-67 staining. **E**, Apoptosis of rPASMCs evaluated by cleaved caspase 3 expression in the hnRNPA1 siRNA-treated group compared with that in the negative control siRNA-treated group by staining for cleaved caspase 3. Orange, yellow, blue, and green colors show normoxia + control siRNA, normoxia + hnRNPA1 siRNA, hypoxia + control siRNA, and hypoxia + hnRNPA1 siRNA, respectively. \**P* < 0.05; \*\**P* < 0.01. Statistical analysis was performed using the Wilcoxon signed-rank test. Values are the mean ± SE, n = 6 per group. CCK, cell counting kit; hnRNPA1, heterogeneous nuclear ribonucleoprotein A1; rPASMC, rat pulmonary artery smooth muscle cell; siRNA, small interfering RNA.

Automatic capillary western blotting analysis by Simple Western (Figure 4A–4D) showed that in normal rPASMCs, hnRNPA1 was firmly suppressed by hnRNPA1 siRNA; the expression of the scaffold protein IQ motif containing GTPase activating protein 1 (IQGAP1), which is reportedly associated with hnRNPA1^21^, was slightly reduced; and the expression of PKM2, which is reportedly involved in cell proliferation, was also reduced. The higher proliferative capacity of hypoxia-treated rPASMCs than that of normal rPASMCs could be associated with elevated hnRNPA1 and PKM2 expression. Knockdown of hnRNAP1 in hypoxia-treated rPASMCs also resulted in markedly decreased hnRNPA1 and PKM2 expression and cell proliferation.

**Figure 4.**
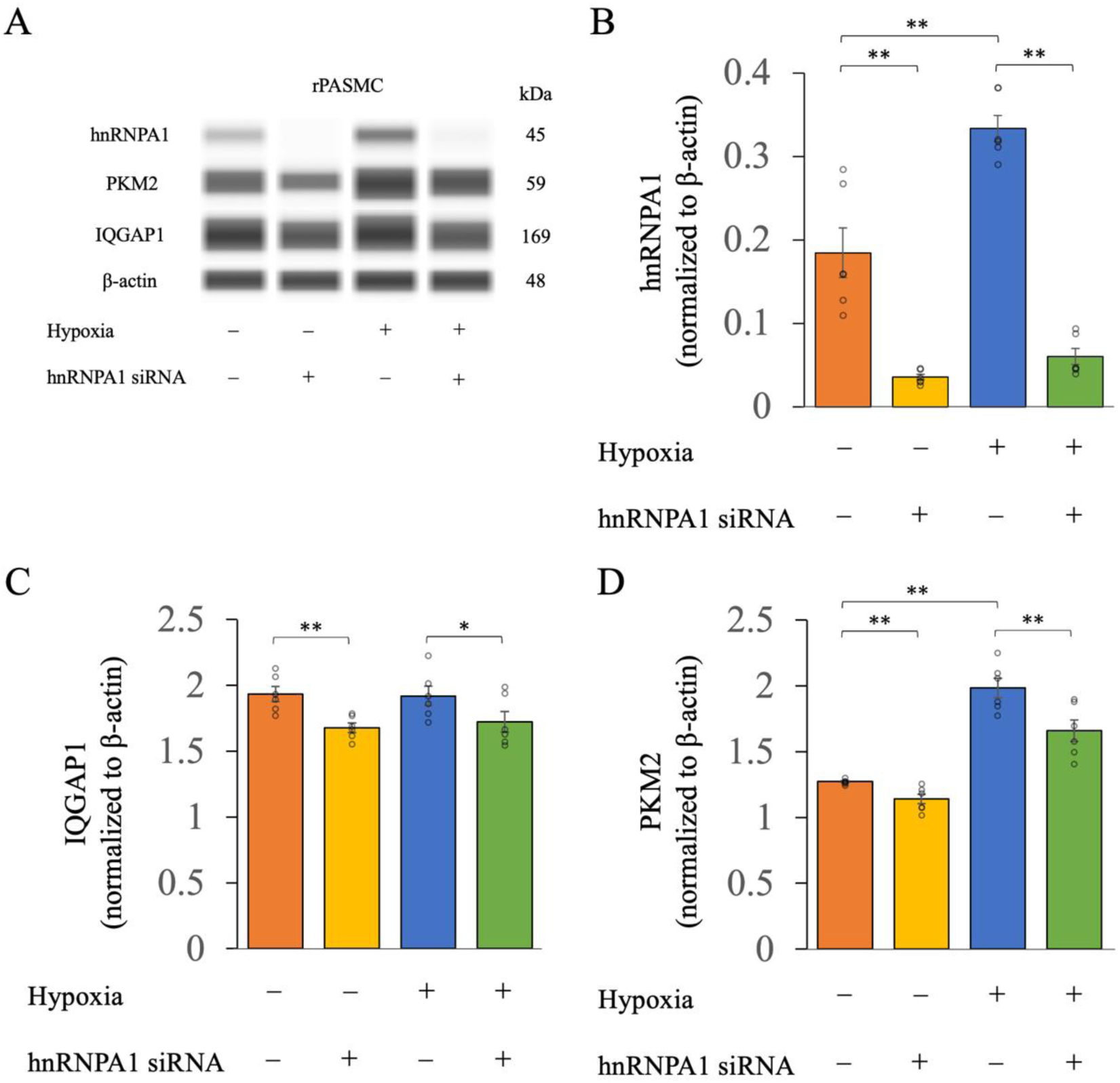
Effect of hnRNPA1 siRNA on protein expression in normal rPASMCs and hypoxia-treated rPASMCs. **A**, hnRNPA1, IQGAP1, PKM2 and β-actin protein expression in normal rPASMCs and hypoxia-treated rPASMCs treated with negative control siRNA or hnRNPA1 siRNA. **B–D**, Ratio of hnRNPA1/β-actin **(B)** PKM2/β-actin **(C)** and IQGAP1/β-actin **(D)** in normal rPASMCs and hypoxia-treated rPASMCs treated with negative control or hnRNPA1 siRNAs. Orange, yellow, blue, and green colors show normoxia + control siRNA, normoxia + hnRNPA1 siRNA, hypoxia + control siRNA, and hypoxia + hnRNPA1 siRNA, respectively. \**P* < 0.05; \*\**P* < 0.01. Statistical analysis was performed using the Wilcoxon signed-rank test. Values are the mean ± SE, n = 6 per group. hnRNPA1, heterogeneous nuclear ribonucleoprotein A1; IQGAP1, IQ motif containing GTPase activating protein 1; PKM2, pyruvate kinase M 2; rPASMC, rat pulmonary artery smooth muscle cell; siRNA, small interfering RNA.

### Inhibition of hnRNPA1 in rPASMCs using hnRNPA1 Inhibitor

In hypoxia-treated rPASMCs, inhibition of hnRNPA1 with tetracaine significantly reduced hnRNPA1 levels in a concentration-dependent manner and reduced cell proliferative capacity (Figure 5A–5E); a similar result has been previously reported for malignant mouse (B16) and human (A375) melanoma cell lines.^19^

**Figure 5.**
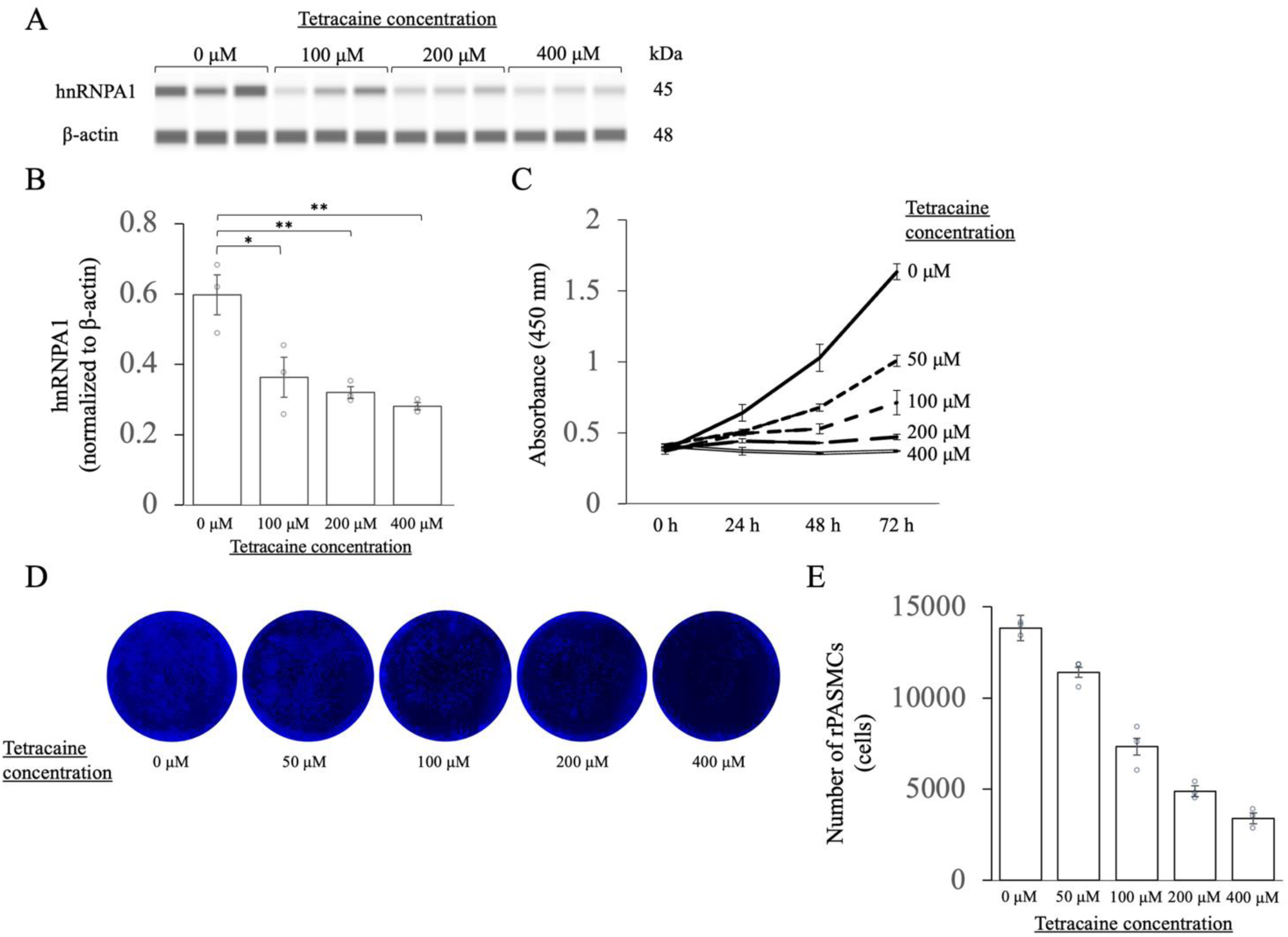
Effect of administration of hnRNPA1 inhibitor to hypoxia-treated rPASMCs on hnRNPA1 expression and cell proliferative capacity. **A**, hnRNPA1 and β-actin protein expression in hypoxia-treated rPASMCs treated with tetracaine. **B**, Ratio of hnRNPA1/β-actin in hypoxia-treated rPASMCs treated with tetracaine. **C**, Proliferative capacity of hypoxia-treated rPASMCs treated with tetracaine, measured via the CCK-8 assay. **D**, rPSAMCs stained with Hoechst dye 72 h after tetracaine treatment. **E**, Number of rPASMCs adhering to the bottom of 96-well plates 72 h after tetracaine treatment. \**P* < 0.05; \*\**P* < 0.01. Statistical analysis was performed using the Wilcoxon signed-rank test. Values are the mean ± SE, n = 3 per group. CCK, cell counting kit; hnRNPA1, heterogeneous nuclear ribonucleoprotein A1; rPASMC, rat pulmonary artery smooth muscle cell.

### Inhibition of hnRNPA1 in SuHx Rats using an hnRNPA1 Inhibitor

Upon administration of tetracaine to SuHx rats at 8 weeks after SU5416 injection, when PLs were more likely to form (Figure S2), a reduction in RVSP was observed in the treated group (82.8 ± 4.1 mmHg, *P* < 0.05) compared with that in the non-treated group (101.3 ± 7.6 mmHg; Figure 6A). Additionally, Fulton’s index decreased in the treated group (0.50 ± 0.03, *P* < 0.01) relative to its value in the non-treated group (0.62 ± 0.01; Figure 6B).

**Figure 6.**
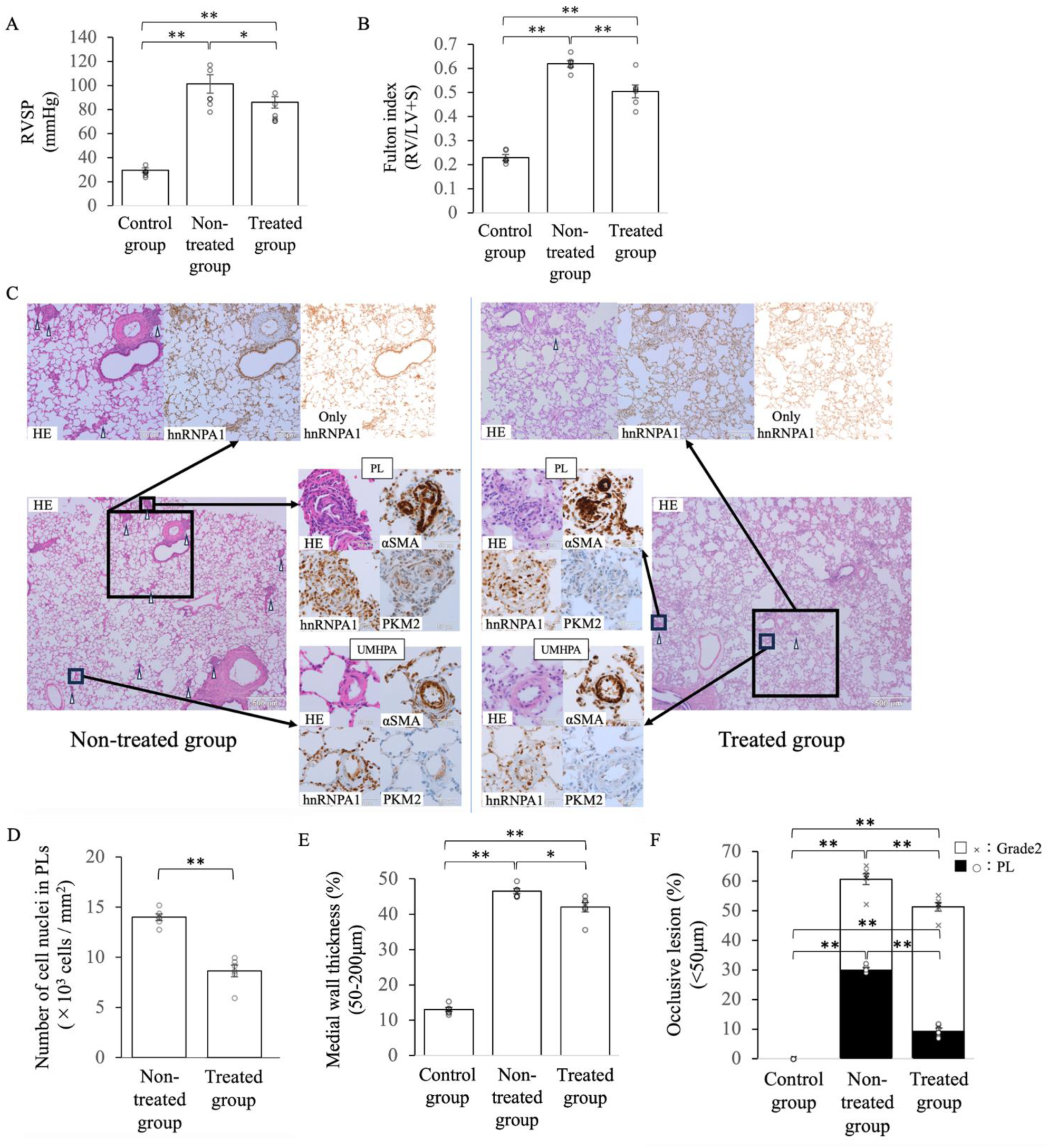
Effects of hnRNPA1 inhibitor on SuHx rats. **A**, Right ventricular systolic pressure (RVSP). **B**, Fulton’s index, weight of the right ventricle to the left ventricle + septum (RV/LV+S). **C**, Comparison of pathology and immunohistochemistry between non-treated and tetracaine-treated groups (scale bar = 500 μm; large inset, scale bar = 200 μm; small inset, scale bar = 50 μm). For hnRNPA1 images, diaminobenzidene and hematoxylin staining are removed by ImageJ, and only hnRNPA1 staining is detected. In HE staining, white arrow heads indicate PLs. **D**, Total number of cell nuclei in PLs (×10^3^ cells/mm^2^) obtained by ImageJ. **E**, Percentage of medial wall thickness. **F**, Percentage of occlusive lesions among the vessels. \**P* < 0.05; \*\**P* < 0.01. Statistical analysis was performed using the Wilcoxon signed-rank test. Values are the mean ± SE, n = 6 per group. αSMA, α-smooth muscle actin; HE, hematoxylin and eosin; hnRNPA1, heterogeneous nuclear ribonucleoprotein A1; PKM2; pyruvate kinase M2; PL, plexiform lesion; SuHx; SU5416 combined with hypoxia; UMHPA, unobstructed pulmonary arteries with medial hypertrophy.

Relative to the level in the non-treated group (66.8 ± 16.3 ng/mL), serum hnRNPA1 concentration was also suppressed in the treated group (21.7 ± 3.3 ng/mL, *P <* 0.05; Figure S6A). Figure 6C shows pathological effects of the hnRNPA1 inhibitor on SuHx rats. hnRNPA1 staining levels were reduced in the treated group and PKM2 levels also tended to decrease. The degree of hnRNPA1 immunostaining (mean gray value), as assessed via ImageJ, was reduced in the treated group compared with that in the non-treated group (Figure S6B). Therefore, tetracaine suppresses hnRNPA1 expression *in vivo*.

The number of cell nuclei in PLs was reduced in the treated group compared with that in the non-treated group (Figure 6D). In the treated group, there were lesser PAs with thickened medial walls (42.0 ± 1.4%, *P* < 0.05) than in the non-treated group (46.5 ± 0.7%; Figure 6E). Figure 6C show PLs (white arrowheads), which were visibly decreased in the treated group. Fewer PLs (9.5 ± 0.8%, *P* < 0.01) were present in the treated group than in the non-treated group (30.0 ± 0.6%; Figure 6F).

In summary, we observed a reduced medial wall thickness, vascular occlusion rate, RVSP, and Fulton’s index in the treated group; pathological evaluation showed a trend toward fewer PLs in the treated group than in the non-treated group, with a reduced number of cell nuclei per PL.

## DISCUSSION

For the first time, DEPs involved in PA remodeling in PAH and PLs were explored via LCM–MS. The main findings of our study were as follows: 1) hnRNPA1 was among the DEPs, which was shown to be expressed in rPASMCs and involved in their proliferation, and 2) suppression of hnRNPA1 reduced PA remodeling in PAH and PLs *in vivo*.

In previous studies, LCM–MS was applied to fresh,^22^ flash-frozen,^23^ and FFPE tissues.^7, 8^ We first experimented on frozen sections but had difficulty in visually identifying the small UMHPAs and obtaining the necessary amount of protein for analysis. Although the protein detection rate is higher when fresh or flash-frozen tissues are used instead of FFPE tissues,^22, 23^ FFPE sections were selected for use because the luminal structure is maintained, which can be easily viewed in the sections. Despite the disadvantage of lower protein detection, we detected protein amounts similar to those previously reported for FFPE tissues^7, 8, 15, 16^ and, following previously published methods, were able to effectively analyze the FFPE tissue using LCM–MS.

To the best of our knowledge, this is the first study to directly examine DEPs in UMHPAs and PLs using LCM–MS. Few LCM–MS studies have previously targeted pulmonary vessels; however, this is the first report examining PLs.^7, 8^ A literature search was conducted to understand the functions of the DEPs detected by our proteomic analysis. Many retrieved proteins are known and reportedly associated with PA remodeling in PAH. In UMHPAs, levels of ANXA3, CAV1, myeloid-associated differentiation marker, TAGLN, and transforming growth factor beta-1-induced transcript 1 protein, which are shown to be associated with PA remodeling in PAH,^9, 24–28^ were elevated. In PLs, in addition to elevated levels of C3, FABP5, and MIF, which are reportedly associated with PA remodeling in PAH, and levels of TMSB4 and PRDX2, which are reportedly protective against PA remodeling in PAH, were elevated.^29–32^ Thus, even in PAH lesions, elevated levels of proteins varied according to lesion status. Moreover, we identified several DEPs, including hnRNPA1, whose role in PA remodeling in PAH has yet to be established. A similar approach has been reported using transcriptome analysis to compare gene expression in PLs and UMHPAs, in which C3, CNN1, TAGLN, TMSB4, MGP, myosin heavy chain 11 and MYL9 exhibited similar behavior to that observed in our study, but not all DEPs were matched.^10^ Immunostaining for several DEPs were in line with those obtained from our proteome analysis. The differences between the results of proteomic analysis and transcriptome analysis can be attributed to species differences and the inherent discrepancy between gene and protein expression levels, which may not always align perfectly.^33^

Among the pathways enhanced in UMHPAs in the enrichment analyses, “cell–extracellular matrix interactions,” “focal adhesion,” “bacterial invasion of epithelial cells,” and “regulation of smooth muscle cell proliferation” are related to PA remodeling in PAH.^34–37^ Similarly, among the pathways enhanced in PLs, “innate immune system,” “G2/M checkpoints,” and “cellular response to reactive oxygen species” are related to PA remodeling in PAH.^38–40^ Thus, enhancement of different pathways in UMHPAs and PLs reflect the involvement of DEPs in these pathological conditions. This indicates that therapeutic targets may differ depending on the disease status and severity.

Based on our results, we focused on hnRNPA1, an RBP that has recently attracted attention for its relationship with PA remodeling in PAH.^20^ Polypyrimidine tract binding protein 1, hnRNPA2/B1, and nucleoli are RBPs strongly associated with PA remodeling in PAH.^20, 41–44^ In arteriosclerosis, hnRNPA1 inhibits the proliferative capacity of systemic vascular smooth muscle cells.^21^ In contrast, hnRNPA1 promotes the proliferative capacity of PASMCs in patients with chronic thromboembolic pulmonary hypertension and its level is increased in patients with hypoxia and PAH.^20, 45^ Thus, hnRNPA1 exhibits dual effects, and its involvement in PA remodeling in PAH remains to be elucidated. The role of hnRNPA1 is also of interest in cancer growth, as it reportedly enhances the proliferation and migratory ability of undifferentiated cells by increasing PKM2 expression because hnRNPA1 can facilitate the switch from PKM1 to PKM2.^46^ Based on these contradictory results, we hypothesized that hnRNPA1 inhibits the proliferative capacity of mature differentiated PASMCs but enhances the proliferative capacity of cancer-like undifferentiated PASMCs.

In mature vascular smooth muscle cells, hnRNPA1 siRNA transfection increases the expression of scaffold protein IQGAP1, enhances proliferative capacity, and enables atherosclerosis progress.^21^ We hypothesized that hnRNPA1 suppresses the proliferative capacity of differentiated cells and performed knockdown experiments in normal and hypoxia-treated rPASMCs. Following siRNA transfection of normal rPASMCs, the expression of IQGAP1 and PKM2 as well as the proliferative capacity were reduced. Hypoxia-treated rPASMCs exhibited similar features. Therefore, siRNA transfection reduced the proliferative capacity in rPASMC under both conditions, indicating that their kinetics were different from those of systemic VSMCs.

Nusilusen, a clinically applied drug for the treatment of spinal muscular atrophy, is a potent inhibitor of hnRNPA1 but is expensive, and the concentrations for use in the experimental model have not been determined.^13^ Instead, we used tetracaine, an inexpensive local anesthetic, to suppresses hnRNPA1 in rPASMCs and SuHx rats, as previously demonstrated in an experimental melanoma model,^19^ and investigated the changes in PA remodeling in PAH.

Suppressing hnRNPA1 with tetracaine reduced the proliferative capacity of hypoxia-treated rPASMCs and inhibited PLs and PA remodeling in PAH in SuHx rats. Importantly, the results of cellular and animal experiments were consistent. Based on these results, we inferred that hnRNPA1 is strongly associated with PA remodeling in PAH and PLs, similar to a previous report on chronic thromboembolic pulmonary hypertension.^45^ Suppressing hnRNPA1 in the early stages of PAH may be associated with a low risk of worsening PA remodeling in PAH, although the risk of atherosclerosis in the systemic vessels should be considered. Results of previous reports and the present study provide evidence that similar to other RBPs, suppressing the excessive increase in hnRNPA1 in undifferentiated PASMCs and immature PLs may suppress PA remodeling in PAH and PLs.^20, 41–44^

The strengths of this study are as follows. For the first time, we comprehensively examined the direct proteomic remodeling in PLs by LCM–MS. We identified several DEPs, prior to and following PL formation, with unclear relevance to PA remodeling in PAH, which were not detected by previously reported transcriptome analysis. There were significant differences in protein and pathway enrichment between UMHPAs and PLs, which may lead to the identification of different therapeutic targets. In addition, hnRNPA1 is strongly expressed in PLs, and its suppression in normal and hypoxia-treated rPASMCs, and SuHx rats showed that hnRNPA1 might be strongly associated with PA remodeling in PAH and PLs. Thus, hnRNPA1 is a potentially useful target for the development of therapies for severe PAH.

In this study, we used hypoxia-treated rPASMCs, whose results may differ from those obtained had we used primary PASMCs from rats with PAH. In addition, we used the general anesthetic tetracaine to suppress hnRNPA1. Therefore, the possibility exists that respiratory depression due to the sedative effects, or other effects of the drug, affect PA remodeling in PAH. However, none of the rats appeared to be sedated or have respiratory depression after drug administration; instead, they all moved normally, suggesting that the sedative effect of the drug did not have any substantial impact at the administered dose. Third, functional analysis of other DEPs is underway, and it is possible that other important DEPs are present. Finally, this was a functional analysis of hnRNPA1 in rats, and the results may differ from those in humans. Further studies in humans are needed, but as samples obtained from humans present a range of clinical variables such as disease stage and medication type, it is possible that studies in a uniform animal model may have detected important DEPs.

In conclusion, we detected several DEPs that could be related to PA remodeling in PAH and PLs and potential therapeutic targets in PAH. hnRNPA1 may play a prominent role in PA remodeling in PAH and PLs, and further studies are warranted to validate our findings before its utility as a therapeutic target or biomarker can be firmly established. There are currently few drugs that have a therapeutic effect on the PLs in severe PAH. Our research has elucidated some of the mechanisms involved in PA remodeling in PAH and PLs, which may be useful in the treatment of PLs.

## Non-Standard Abbreviations and Acronyms

αSMA: α-smooth muscle actin
ANXA3: annexin A3
ACTB: β-actin
CALML3: calmodulin-like protein 3
CNN1: calponin 1
CAV1: caveolin 1
CCK-8: cell counting kit-8
C3: complement C3
C4: complement C4
DEP: differentially expressed protein
FABP5: fatty acid-binding protein 5
FC: fold-change
FFPE: formalin-fixed paraffin-embedded
hnRNPA1: heterogeneous nuclear ribonucleoprotein A1
IQGAP1: IQ motif containing GTPase activating protein 1
LCM–MS: laser-capture microdissection coupled with mass spectrometry
LCA: leukocyte common antigen
LOQ: limit of quantification
MGP: matrix Gla protein
MEM: minimum essential medium
MIF: migration inhibitory factor
MYL9: myosin light chain 9
PRDX2: peroxiredoxin 2
PBS: phosphate-buffered saline
PECAM-1: platelet endothelial cell adhesion molecule 1
PL: plexiform lesion
PAH: pulmonary arterial hypertension
PA: pulmonary artery
PKM2: pyruvate kinase 2
RBP: RNA-binding protein
rPASMCs: rat pulmonary artery smooth muscle cells
siRNA: small interfering RNA
SuHx: SU5416 combined with hypoxia
TAGLN: transgelin
TMSB4: thymosin beta 4
UMHPAs: unobstructed pulmonary arteries with medial hypertrophy

## ARTICLE INFORMATION

## Acknowledgments

The authors wish to acknowledge Kentaro Taki and the Division for Medical Research Engineering, Nagoya University Graduate School of Medicine, for technical support provided with Cytation, usage of the LMD 7000 and CentriVap, and liquid chromatography–tandem mass spectrometry analysis. The authors are grateful to Azusa Okamoto for the skillful technical assistance. The authors would also like to thank Editage (https://www.Editage.jp) for English language editing.

## Supplemental Material

Supplemental Methods

Figures S1–S6

Tables S1–S2

Major Resources Table

## Study Specific Approval

All animals received humane care, and all experimental procedures were approved by the Animal Care and Use Committee of Nagoya University (M240320-001 and M240315-001). In addition, all procedures conformed to the guidelines from Directive 2010/63/EU of the European Parliament on the protection of animals used for scientific purposes.

## REFERENCES

1. Humbert M, Lau EMT, Montani D, Jaïs X, Sitbon O, Simonneau G. Advances in therapeutic interventions for patients with pulmonary arterial hypertension. Circulation. 2014;130:2189–2208. doi:10.1161/CIRCULATIONAHA.114.006974

2. Jonigk D, Golpon H, Bockmeyer CL, Maegel L, Hoeper MM, Gottlieb J, Nickel N, Hussein K, Maus U, Lehmann U, et al. Plexiform lesions in pulmonary arterial hypertension composition, architecture, and microenvironment. Am J Pathol. 2011;179:167–179. doi:10.1016/j.ajpath.2011.03.040

3. Cascino TM, Sahay S, Moles VM, McLaughlin VV. A new day has come: sotatercept for the treatment of pulmonary arterial hypertension. J Heart Lung Transplant. 2025;44:1–10. doi:10.1016/j.healun.2024.09.021

4. Espina V, Wulfkuhle JD, Calvert VS, VanMeter A, Zhou W, Coukos G, Geho DH, Petricoin EF 3rd, Liotta LA. Laser-capture microdissection. Nat Protoc. 2006;1:586–603. doi:10.1038/nprot.2006.85

5. Xu BJ. Combining laser capture microdissection and proteomics: methodologies and clinical applications. Proteomics Clin Appl. 2010;4:116–123. doi:10.1002/prca.200900138

6. Dilillo M, Pellegrini D, Ait-Belkacem R, de Graaf EL, Caleo M, McDonnell LA. Mass spectrometry imaging, laser capture microdissection, and LC–MS/MS of the same tissue section. J Proteome Res. 2017;16:2993–3001. doi:10.1021/acs.jproteome.7b00284

7. Fayyaz AU, Sabbah MS, Dasari S, Griffiths LG, DuBrock HM, Wang Y, Charlesworth MC, Borlaug BA, Jenkins SM, Edwards WD, Redfield MM. Histologic and proteomic remodeling of the pulmonary veins and arteries in a porcine model of chronic pulmonary venous hypertension. Cardiovasc Res. 2023;119:268–282. doi:10.1093/cvr/cvac005

8. Herrera JA, Mallikarjun V, Rosini S, Montero MA, Lawless C, Warwood S, O’Cualain R, Knight D, Schwartz MA, Swift J. Laser capture microdissection coupled mass spectrometry (LCM-MS) for spatially resolved analysis of formalin-fixed and stained human lung tissues. Clin Proteomics. 2020;17:24. doi:10.1186/s12014-020-09287-6

9. Abdul-Salam VB, Wharton J, Cupitt J, Berryman M, Edwards RJ, Wilkins MR. Proteomic analysis of lung tissues from patients with pulmonary arterial hypertension. Circulation. 2010;122:2058–2067. doi:10.1161/CIRCULATIONAHA.110.972745

10. Tuder RM, Gandjeva A, Williams S, Kumar S, Kheyfets VO, Hatton-Jones KM, Starr JR, Yun J, Hong J, West NP, et al. Digital spatial profiling identifies distinct molecular signatures of vascular lesions in pulmonary arterial hypertension. Am J Respir Crit Care Med. 2024;210:329–342. doi:10.1164/rccm.202307-1310OC

11. Abe K, Toba M, Alzoubi A, Ito M, Fagan KA, Cool CD, Voelkel NF, McMurtry IF, Oka M. Formation of plexiform lesions in experimental severe pulmonary arterial hypertension. Circulation. 2010;121:2747–2754. doi:10.1161/CIRCULATIONAHA.109.927681

12. Shinohara T, Sawada H, Otsuki S, Yodoya N, Kato T, Ohashi H, Zhang E, Saitoh S, Shimpo H, Maruyama K, et al. Macitentan reverses early obstructive pulmonary vasculopathy in rats: early intervention in overcoming the survivin-mediated resistance to apoptosis. Am J Physiol Lung Cell Mol Physiol. 2015;308:L523–L538. doi:10.1152/ajplung.00129.2014

13. Reilly A, Chehade L, Kothary R. Curing SMA: are we there yet? Gene Ther. 2023;30:8–17. doi:10.1038/s41434-022-00349-y

14. Otsuki S, Sawada H, Yodoya N, Shinohara T, Kato T, Ohashi H, Zhang E, Imanaka-Yoshida K, Shimpo H, Maruyama K, et al. Potential contribution of phenotypically modulated smooth muscle cells and related inflammation in the development of experimental obstructive pulmonary vasculopathy in rats. PLoS One. 2015;10:e0118655. doi:10.1371/journal.pone.0118655

15. Drummond ES, Nayak S, Ueberheide B, Wisniewski T. Proteomic analysis of neurons microdissected from formalin-fixed, paraffin-embedded Alzheimer’s disease brain tissue. Sci Rep. 2015;5:15456. doi:10.1038/srep15456

16. Drummond E, Nayak S, Pires G, Ueberheide B, Wisniewski T. Isolation of amyloid plaques and neurofibrillary tangles from archived Alzheimer’s disease tissue using laser-capture microdissection for downstream proteomics. Methods Mol Biol. 2018;1723:319–334. doi:10.1007/978-1-4939-7558-7_18

17. Cotto-Rios XM, Békés M, Chapman J, Ueberheide B, Huang TT. Deubiquitinases as a signaling target of oxidative stress. Cell Rep. 2012;2:1475–1484. doi:10.1016/j.celrep.2012.11.011

18. Wei S, Lin L, Jiang W, Chen J, Gong G, Sui D. Naked cuticle homolog 1 prevents mouse pulmonary arterial hypertension via inhibition of Wnt/β-catenin and oxidative stress. Aging (Albany NY). 2023;15:11114–11130. doi:10.18632/aging.205105

19. Huang X, Chen Y, Yi J, Yi P, Jia J, Liao Y, Feng J, Jiang X. Tetracaine hydrochloride induces cell cycle arrest in melanoma by downregulating hnRNPA1. Toxicol Appl Pharmacol. 2022;434:115810. doi:10.1016/j.taap.2021.115810

20. Zhang H, Brown RD, Stenmark KR, Hu CJ. RNA-binding proteins in pulmonary hypertension. Int J Mol Sci. 2020;21:3757. doi:10.3390/ijms21113757

21. Zhang L, Chen Q, An W, Yang F, Maguire EM, Chen D, Zhang C, Wen G, Yang M, Dai B, et al. Novel pathological role of hnRNPA1 (heterogeneous nuclear ribonucleoprotein A1) in vascular smooth muscle cell function and neointima hyperplasia. Arterioscler Thromb Vasc Biol. 2017;37:2182–2194. doi:10.1161/ATVBAHA.117.310020

22. Li C, Hong Y, Tan YX, Zhou H, Ai JH, Li SJ, Zhang L, Xia QC, Wu JR, Wang HY, et al. Accurate qualitative and quantitative proteomic analysis of clinical hepatocellular carcinoma using laser capture microdissection coupled with isotope-coded affinity tag and two-dimensional liquid chromatography mass spectrometry. Mol Cell Proteomics. 2004;3:399–409. doi:10.1074/mcp.M300133-MCP200

23. Clair G, Piehowski PD, Nicola T, Kitzmiller JA, Huang EL, Zink EM, Sontag RL, Orton DJ, Moore RJ, Carson JP, et al. Spatially-resolved proteomics: rapid quantitative analysis of laser capture microdissected alveolar tissue samples. Sci Rep. 2016;6:39223. doi:10.1038/srep39223

24. Park JE, Lee DH, Lee JA, Park SG, Kim NS, Park BC, Cho S. Annexin A3 is a potential angiogenic mediator. Biochem Biophys Res Commun. 2005;337:1283–1287. doi:10.1016/j.bbrc.2005.10.004

25. Mathew R. Critical role of caveolin-1 loss/dysfunction in pulmonary hypertension. Med Sci (Basel). 2021;9:58. doi:10.3390/medsci9040058

26. Zhang R, Shi L, Zhou L, Zhang G, Wu X, Shao F, Ma G, Ying K. Transgelin as a therapeutic target to prevent hypoxic pulmonary hypertension. Am J Physiol Lung Cell Mol Physiol. 2014;306:L574–L583. doi:10.1152/ajplung.00327.2013

27. Sun L, Lin P, Chen Y, Yu H, Ren S, Wang J, Zhao L, Du G. miR-182-3p/Myadm contribute to pulmonary artery hypertension vascular remodeling via a KLF4/p21-dependent mechanism. Theranostics. 2020;10:5581–5599. doi:10.7150/thno.44687

28. Ji Y, Lisabeth EM, Neubig RR. Transforming growth factor β1 increases expression of contractile genes in human pulmonary arterial smooth muscle cells by potentiating sphingosine-1-phosphate signaling. Mol Pharmacol. 2021;100:53–60. doi:10.1124/molpharm.120.000019

29. Lei Q, Yu Z, Li H, Cheng J, Wang Y. Fatty acid-binding protein 5 aggravates pulmonary artery fibrosis in pulmonary hypertension secondary to left heart disease via activating wnt/β-catenin pathway. J Adv Res. 2022;40:197–206. doi:10.1016/j.jare.2021.11.011

30. Jalce G, Guignabert C. Multiple roles of macrophage migration inhibitory factor in pulmonary hypertension. Am J Physiol Lung Cell Mol Physiol. 2022;318:L1–L9. doi:10.1152/ajplung.00234.2019

31. Federti E, Matté A, Ghigo A, Andolfo I, James C, Siciliano A, Leboeuf C, Janin A, Manna F, Choi SY, et al. Peroxiredoxin-2 plays a pivotal role as multimodal cytoprotector in the early phase of pulmonary hypertension. Free Radic Biol Med. 2017;112:376–386. doi:10.1016/j.freeradbiomed.2017.08.004

32. Wei C, Kim IK, Li L, Wu L, Gupta S. Thymosin beta 4 protects mice from monocrotaline-induced pulmonary hypertension and right ventricular hypertrophy. PLoS One. 2021;9:e110598. doi:10.1371/journal.pone.0110598

33. Bathke J, Konzer A, Remes B, McIntosh M, Klug G. Comparative analyses of the variation of the transcriptome and proteome of *Rhodobacter sphaeroides* throughout growth. BMC Genomics. 2019;20:358. doi:10.1186/s12864-019-5749-3

34. Link PA, Farkas L, Heise RL. Using extracellular matrix derived from sugen-chronic hypoxia lung tissue to study pulmonary arterial hypertension. Front Pharmacol. 2023;14:1192798. doi:10.3389/fphar.2023.1192798

35. Wang R, Xu J, Wu J, Gao S, Wang Z. Angiotensin-converting enzyme 2 alleviates pulmonary artery hypertension through inhibition of focal adhesion kinase expression. Exp Ther Med. 2021;22:1165. doi:10.3892/etm.2021.10599

36. Zhang C, Zhang T, Lu W, Duan X, Luo X, Liu S, Chen Y, Li Y, Chen J, Liao J, et al. Altered airway microbiota composition in patients with pulmonary hypertension. Hypertension. 2020;76:1589–1599. doi:10.1161/HYPERTENSIONAHA.120.15025

37. Huetsch JC, Suresh K, Shimoda LA. Regulation of smooth muscle cell proliferation by NADPH oxidases in pulmonary hypertension. Antioxidants (Basel). 2019;8:56. doi:10.3390/antiox8030056

38. Kherbeck N, Tamby MC, Bussone G, Dib H, Perros F, Humbert M, Mouthon L. The role of inflammation and autoimmunity in the pathophysiology of pulmonary arterial hypertension. Clin Rev Allergy Immunol. 2013;44:31–38. doi:10.1007/s12016-011-8265-z

39. Sankhe S, Manousakidi S, Antigny F, Arthur Ataam J, Bentebbal S, Ruchon Y, Lecerf F, Sabourin J, Price L, Fadel E, et al. T-type Ca2+ channels elicit pro-proliferative and anti-apoptotic responses through impaired PP2A/Akt1 signaling in PASMCs from patients with pulmonary arterial hypertension. Biochim Biophys Acta Mol Cell Res. 2017;1864:1631–1641. doi:10.1016/j.bbamcr.2017.06.018

40. Ahmad Y, Sharma NK, Ahmad MF, Sharma M, Garg I, Bhargava K. Proteomic identification of novel differentiation plasma protein markers in hypobaric hypoxia-induced rat model. PLoS One. 2014;9:e98027. doi:10.1371/journal.pone.0098027

41. Zhang H, Wang D, Li M, Plecitá-Hlavatá L, D’Alessandro A, Tauber J, Riddle S, Kumar S, Flockton A, McKeon BA, et al. Metabolic and proliferative state of vascular adventitial fibroblasts in pulmonary hypertension is regulated through a microRNA-124/PTBP1 (polypyrimidine tract binding protein 1)/pyruvate kinase muscle axis. Circulation. 2017;136:2468–2485. doi:10.1161/CIRCULATIONAHA.117.028069

42. Ruffenach G, Medzikovic L, Aryan L, Li M, Eghbali M. HNRNPA2B1: RNA-binding protein that orchestrates smooth muscle cell phenotype in pulmonary arterial hypertension. Circulation. 2022;146:1243–1258. doi:10.1161/CIRCULATIONAHA.122.059591

43. Lee J, Kang H. Nucleolin regulates pulmonary artery smooth muscle cell proliferation under hypoxia by modulating miRNA expression. Cells. 2023;12:817. doi:10.3390/cells12050817

44. Caruso P, Dunmore BJ, Schlosser K, Schoors S, Dos Santos C, Perez-Iratxeta C, Lavoie JR, Zhang H, Long L, Flockton AR, et al. Identification of microRNA-124 as a major regulator of enhanced endothelial cell glycolysis in pulmonary arterial hypertension via PTBP1 (polypyrimidine tract binding protein) and pyruvate kinase M2. Circulation. 2017;136:2451–2467. doi:10.1161/CIRCULATIONAHA.117.028034

45. Liu L, Pang W, Liu J, Xu S, Zhang Z, Hao R, Wan J, Xie W, Tao X, Yang P, et al. Inhibition of heterogeneous nuclear ribonucleoproteins A1 and oxidative stress reduces glycolysis *via* pyruvate kinase M2 in chronic thromboembolic pulmonary hypertension. J Transl Int Med. 2024;12:437–451. doi:10.2478/jtim-2022-0051

46. Zhao B, Lv X, Zhao X, Maimaitiaili S, Zhang Y, Su K, Yu H, Liu C, Qiao T. Tumor-promoting actions of HNRNP A1 in HCC are associated with cell cycle, mitochondrial dynamics, and necroptosis. Int J Mol Sci. 2022;23:10209. doi:10.3390/ijms231810209

